# Electrophysiological and behavioral indices of cognitive conflict processing across adolescence

**DOI:** 10.1101/2020.05.08.084194

**Authors:** Knut Overbye, Kristine B. Walhovd, Anders M. Fjell, Christian K. Tamnes, Rene J. Huster

**Author notes:** Correspondence to: Knut Overbye, Department of Psychology, University of Oslo, PO Box 1094 Blindern, 0317 Oslo, Norway.

## Abstract

Cognitive control enables goal-oriented adaptation to a fast-changing environment and has a slow developmental trajectory that spans into young adulthood. The specifics of this development are still poorly understood, as are the neurodevelopmental mechanisms that drive it. In a cross-sectional sample of participants 8-19 years old (n = 108), we used blind source separation of EEG data recorded in a Flanker task to derive electrophysiological measures of attention and the processing of cognitive conflict, including a frontal negative component corresponding to the N2 and a parietal positive component corresponding to the P3. Additionally, we examined multiple behavioral measures of interference control derived from the Flanker, Stroop, and Anti-saccade tasks. We found a positive association between age and the amplitude of the parietal positive component, while there was no relationship between age and the amplitude of the frontal negative component. A stronger frontal negative amplitude was, however, age-independently related to better performance on both Stroop and Anti-saccade measures of interference control. Finally, we examined post-conflict behavioral adjustment on the Flanker task. A Gratton effect was found with slower reaction times on current congruent and better accuracy on current incongruent trials when preceded by incongruent as opposed to congruent trials. The Gratton effect on accuracy was positively associated with age. Together, the findings suggest a multifaceted developmental pattern in neurocognitive mechanisms for conflict processing across adolescence, with a more protracted development of the parietal positive compared to the frontal negative component.

## INTRODUCTION

The quick and precise resolution of ambiguous, irreconcilable or otherwise conflicting information is a central aspect to adaptive behavior and decision making. The set of cognitive processes that are used to resolve such conflict is known as cognitive control. The cognitive control of thoughts and actions is among the cognitive functions to develop the slowest during ontogeny, showing continued improvement through adolescence and well into young adulthood (Crone & Steinbeis, 2017; Zelazo & Müller, 2002).

Processes of cognitive control are thought to be triggered when there is conflict between competing neural representations (Egner, 2008). In other words, perceived conflicts in information processing serves as a signal to bring processes of cognitive control to bear (Botvinick, Braver, Barch, Carter, & Cohen, 2001). Sources of conflict can range from basic and immediate events such as present distractions to more complex and drawn out processes such as the delay of gratification (Crone & Steinbeis, 2017). From this, it follows that cognitive control is a broad construct encompassing a variety of functions, from fairly basic and immediate to slow processes influenced by long-term planning (Crone & Steinbeis, 2017). In the present study we focus on a relatively basic and immediate variety, specifically stimulus interference control, which describes the ability to overcome distraction from irrelevant information.

Interference control is commonly measured with behavioral tasks such as the Stroop Color-Word Interference Task, the Anti-saccade task, or the Eriksen Flanker task; tests in which ignoring distracting cues is essential for task performance. Performance on these tasks has been shown to improve through adolescence (Fukushima, Hatta, & Fukushima, 2000; Ikeda, Okuzumi, Kokubun, & Haishi, 2011; Klein & Foerster, 2001; Luna, Garver, Urban, Lazar, & Sweeney, 2004; Prencipe et al., 2011; Tamnes et al., 2010b), indicating that interference control has a protracted development. Still, little is known about the neural changes underlying development of interference control. Since the detection of conflicts and their resolution hinges on split-second decisions, electrophysiological measures have proven valuable in studying their underlying mechanisms, due to their high temporal resolution. The association between specific electrophysiological and behavioral measures of cognitive control is however not clear (Downes, Bathelt, & De Haan, 2017).

In EEG recordings, the presentation of a conflict-inducing stimulus is generally followed by a negative event related potential (ERP) on the frontal midline of the scalp and a later positive ERP with a parietal scalp topography (Folstein & Van Petten, 2008; Huster, Enriquez-Geppert, Lavallee, Falkenstein, & Herrmann, 2013). The characteristics of these components depend on the experimental paradigm used. Studies using the Flanker task tend to interpret these ERPs as the N2 and P3, with the N2 occurring fronto-centrally 200-300 ms after stimulus presentation and the P3 appearing roughly 150 ms later, with a centro-parietal topography (Huster et al., 2013). In contrast, studies using the Stroop task, where the ERPs tend to occur later, generally interpret them as the N450 and Conflict Slow Potential (Larson, Clayson, & Clawson, 2014).

The N2 is believed to be generated in the anterior cingulate cortex, as well as in frontal and superior temporal cortices (Huster, Westerhausen, Pantev, & Konrad, 2010). The magnitude of the N2’s peak seems to relate to the degree of conflict elicited by the task, being more negative if the task is more challenging (Folstein & Van Petten, 2008). Whether a higher N2 amplitude is associated with better response inhibition is not clear (Huster et al., 2013). Huster, Enriquez-Geppert, Pantev, and Bruchmann (2014) found a greater N2 amplitude to be associated with greater accuracy on a Go/No-Go task in adults, while Lamm, Zelazo, and Lewis (2006) found a greater relative N2 amplitude to be associated with worse performance on the Stroop and the Iowa Gambling Task in adolescents. According to a recent review by Lo (2018), no previous study has performed an investigation of the developmental changes of the N2 from childhood to adulthood. However, studies in childhood and early adolescence using a Go/No-Go paradigm, tapping response interference, have generally found the amplitude of the N2 to decrease with age (Lo, 2018). There are exceptions to this trend, however. Notably, a recent large longitudinal study on children between 3 and 6 years old found the difference between No-Go and Go amplitudes to remain stable with increasing age (Isbell, Calkins, Cole, Swingler, & Leerkes, 2018). In addition to the N2, there exist several other negative ERPs with a midline frontal scalp topography that are also related to cognitive control. These include the error related negativity (ERN), the feedback related negativity and the correct response negativity (Cavanagh & Frank, 2014), as well as the N450 (Larson et al., 2014). Although these ERPs and the N2 are elicited by different events and show somewhat different associations with behavior, they seem to originate from more or less the same source (Gruendler, Ullsperger, & Huster, 2011), and the degree to which these ERPs reflect separate neurocognitive processes is a matter of ongoing debate (Gratton, Cooper, Fabiani, Carter, & Karayanidis, 2018; Larson et al., 2014; Miller & Cohen, 2001).

The P300 is believed to be largely endogenously generated, related to conscious information processing and is thought to be the first major component of controlled attention (Dehaene, Sergent, & Changeux, 2003; Polich, 2007; Segalowitz & Davies, 2004). The source of the P300 in disputed, with possible contributing sources in both frontal, parietal, temporal and parieto-occipital cortical regions (Bocquillon et al., 2011; Friedman, 2003; Halgren et al., 1995; Mahajan & McArthur, 2015; Smith et al., 1990; Wronka, Kaiser, & Coenen, 2012). The P300 is also frequently split into two different subtypes: the P3a, which relates to stimulus driven attention mechanisms during task processing, and the P3b, which relates to attention and subsequent memory processing (Polich, 2007). The P3a tends to occur slightly earlier and more frontally on the scalp compared to the P3b. An extensive review and meta-analysis of developmental studies on the P300 in auditory oddball tasks by van Dinteren, Arns, Jongsma, and Kessels (2014b) concluded that the P300 amplitude increases from childhood to adulthood. This is also what we recently found for a visual oddball task (Overbye, Huster, Walhovd, Fjell, & Tamnes, 2018).

Beyond these ERPs, there is a tendency for trials following conflict-laden trials to be less affected by interference. This is known as the Gratton effect, or alternatively as the congruency or sequence effect (Botvinick et al., 2001; Gratton, Coles, & Donchin, 1992). Botvinick et al. (2001) suggested that the conflict inducing stimuli triggers an augmentation of cognitive control, which carries over into the next trial, reducing the impact of distracting information. This effect has also been observed in children (Erb & Marcovitch, 2018; Erb, Moher, Song, & Sobel, 2018). A recent study on children and adults by Erb and Marcovitch (2018), found that the Gratton effect was greater in mid childhood compared to preadolescence and adulthood, while no difference was found between pre-adolescents and adults.

In the present study, we performed a broad investigation of age-related differences in cognitive conflict processing across adolescence by examining independent electrophysiological components reflecting the N2 and P3, as well as performance measures from behavioral tasks measuring interference control. Based on previous developmental findings, we expected the N2 amplitude to either decrease or remain stable with age (Isbell et al., 2018; Lo, 2018), while we expected the P3 amplitude to increase with age (van Dinteren, Arns, Jongsma, & Kessels, 2014a; van Dinteren et al., 2014b). We also hypothesized that individual differences in the strength of these components would be associated with performance on tasks reflecting interference control. We predict that a stronger P3 would be associated with better performance. Though the relationship between N2 strength and interference control is debated, we tentatively predict a stronger N2 to be associated with better performance.

## MATERIALS AND METHODS

### Participants

The study was approved by the Norwegian Regional Committee for Medical and Health Research Ethics. Participants between 8 and 19 years of age were recruited to the research project *Neurocognitive Development* (Østby et al., 2009; Tamnes et al., 2010a) through newspaper advertisements, and local schools and workplaces. Written informed consent was provided by all participants over 12 years, and from a parent or guardian of participants younger than 18 years. Oral informed assent was obtained from participants younger than 12 years. Participants aged 16 years or older and a parent completed standardized health interviews regarding each participant. Exclusion criteria included premature birth (<37 weeks of gestation), a history of brain injury or disease, ongoing treatment for a mental disorder, use of psychoactive drugs, and MRI contraindications. Participants were required to be right-handed, fluent Norwegian speakers, and to have normal or corrected-to-normal hearing and vision. A total of 113 children and adolescents fulfilled these criteria and were after MRI scanning deemed free of significant brain injuries or conditions by a neuroradiologist. Five participants were excluded due to task-performance criteria, as described below. Further, eleven more participants were excluded from the EEG analyses due to noisy or incomplete data. Age, sex and estimated IQ for each subsample are reported in **Table 1**. The four-subtest form of the Wechsler Abbreviated Scale of Intelligence was used to estimate IQ (Wechsler, 1999). Participants performed the Stroop Color-Word Interference Task and an Anti-saccade task, as well as a Flanker task during EEG recording. One participant did not perform the Anti-Saccade task and was excluded from analyses using this task. Associations between behavioral performance on the Stroop and Antisaccade tasks and age and cortical thickness have been previously reported (Tamnes et al., 2010b).

**Table 1.** Demographics for the different subsamples used for analyses using only behavioral data and using both behavioral and EEG data.

| Sample descriptives |  |  |
| --- | --- | --- |
|  | Valid behavior | Valid behavior and EEG |
| N | 108 | 94 |
| Age mean (SD) | 14.1 (3.4) | 14.2 (3.4) |
| Age range | 8.3-19.7 | 8.3-19.7 |
| Sex | 56 f / 52 m | 48 f / 46 m |
| Estimated IQ mean (SD) | 109.4 (10.2) | 109.4 (10.3) |

### Flanker Task

Stimuli were presented on a 19-inch computer screen at a viewing distance of approximately 80 cm. Task administration and behavioral recording were done using E-prime software and a Psychology Software Tools Serial Response Box. EEG was recorded during a modified and speeded version of the Eriksen arrow Flanker task (Eriksen & Eriksen, 1974), as previously described elsewhere (Overbye et al., 2019; Tamnes, Fjell, Westlye, Ostby, & Walhovd, 2012). Briefly, stimuli were 2.5° vertical stacks of five 1 cm long white arrows, presented on a black background. Each trial started with a fixation cross presented for a random interval between 1200 and 1800 ms. The interval was randomized to reduce noise from slow-wave activity as well as to increase task difficulty by hampering preparatory behavior. After the fixation cross, four arrows were presented for 80 ms before the target arrow was presented in the middle, together with the flanker arrows, for 60ms. This was done to make the task more difficult by priming for the prepotent response. Finally, a black screen was presented for up to 1440 ms. A total of 416 trials were presented, half of which were congruent, with the target arrow pointing in the same direction as the flankers, and the other half incongruent, with the target and flanker arrows pointing in opposing directions. Left and right pointing target arrows were equally frequent within each condition. Trials were presented in semi-random order, with incongruent trials never appearing more than three times in a row. Participants were asked to respond by pressing one of two buttons depending on the direction of the target arrow. They were also asked to emphasize both speed and accuracy when responding. An individual response time threshold was set for each participant based on their average reaction time on the first 20 trials. If participants responded slower than this threshold on three subsequent trials, they were asked to respond faster through a 1 second text prompt. This was done to increase the error rate. There was a short break half-way through the task. Before the task, participants completed two practice blocks of 12 trials each. In the first of these, both the flankers and the target arrow were presented for slightly longer (150 and 90 ms, respectively), whereas in the second block, the times were the same as in the main task. The Flanker task measures cognitive control and selective attention (Larson et al., 2014; McDermott, Wiesman, Proskovec, Heinrichs-Graham, & Wilson, 2017). We used the following two behavioral inclusion criteria: First, participants were required to have ≥60% accuracy on congruent trials. Second, participants needed to show a significant congruency effect, with reaction times on correct incongruent trials significantly slower than on correct congruent trials as determined by a paired samples t-test (p < .05). These criteria were used to exclude those with suboptimal motivation, or those who did not pay attention to the task or who reponded at random. Three participants were excluded due to low accuracy and two more due to a lack of a congruency effect, yielding a sample of 108 for the behavioral analyses. The median reaction time was calculated separately for correct and incorrect congruent and incongruent trials for each participant. Mean accuracy was calculated separately for congruent and incongruent trials for each participant.

### Color-Word Interference Task

Similar to the Flanker task, the Stroop Color-Word task (Stroop, 1935) is a measure of selective attention and cognitive control. The Stroop task employed was the D-KEFS Color-Word Interference Task (Delis, Kaplan, & Kramer, 2001). For the present study, we were only interested in the abilities uniquely measured by the Incongruent color-word interference condition, with performance on the congruent Color naming condition being regressed out. In the Color naming condition, participants were presented with a sheet of paper with five rows of ten squares, printed in one of three colors (red, blue or green), and were instructed to name the colors one-by-one and row-by-row as fast as possible until finished. In the Incongruent condition, participants were presented with five rows of ten color words (“red”, “blue” and “green”), printed in incongruent colors (red, blue or green), and were asked to name the print colors as fast as possible until finished. Participants were thus required to overcome an overlearned verbal response, i.e. reading the printed words, in order to generate a conflicting response of naming the incongruent ink colors in which the words were printed. Completion time for each condition was measured with a stopwatch. Before each condition, participants practiced on a small number of items. The raw score on the color naming condition was regressed out of the score of each participant’s raw score on the Interference condition to get a purer measure of cognitive control. A ratio measure was also calculated in order to compare our results with previous studies using this approach. This ratio was calculated as Stroop Color Naming reaction time minus Strop Color-Word Interference reaction time, divided by Stroop Color Naming reaction time. Ratio measures do not properly control for the denominator variable (Atchley, Gaskins, & Anderson, 1976), so analyses using this measure are more difficult to interpret than ones using a regression approach.

### Anti-saccade Task

The Anti-saccade task is a measure of inhibition, adapted by Miyake et al. (2000) from Roberts, Hager, and Heron (1994). In each trial, a fixation point was first presented in the middle of the computer screen for a variable duration (randomly selected from a list of nine values in 250 ms intervals between 100 and 2100 ms). A visual cue, a small black square, was then presented on one side of the screen (left or right) for 225 ms, followed by the presentation of a small black target arrow on the opposite side of the screen for 140 ms. The target arrow was then masked by gray cross-hatching for 200 ms. Participants were asked to indicate the direction of the arrow (left, up, or right) with a button press. A Psychology Software Tools Serial Response Box was used. Since the target arrow was presented only briefly before being masked, participants were required to inhibit the reflexive response of looking at the initial cue because doing so would make it more difficult to correctly identify the direction of the target arrow. The task consisted of 18 practice trials and two blocks of 60 target trials each, for a total of 120 target trials. The percentage of target trials answered correctly was the measure of interest.

### EEG Acquisition

Participants performed the Flanker task in an electrically shielded room while seated in a comfortable high-back chair. The electrophysiological recordings were done using 128 EEG channels with an electrode placement based on the 10% system (EasyCap Montage No. 15, http://www.easycap.de/). The sampling rate during recording was set to 1000 Hz. The electrodes used were EasyCap active ringel ectrodes (Ag/AgCl) with impedance conversion circuits integrated into the electrode housing that allows high quality recordings even with high impedance values, thus reducing preparation time and noise. The signals were amplified via a Neuroscan SynAmps2 system and filtered online with a 40 Hz low-pass and a 0.15 Hz high-pass analog filter prior to digitization and saving of the continuous data set. During recording, all electrodes were referenced to an electrode placed on the left mastoid. Eye blinks were recorded with one electrode above and one electrode below the left eye, and a ground electrode was placed anteriorly on the midline.

### EEG Processing

Data pre-processing was done using Matlab and EEGLab. Bad channels, with insufficient or corrupt data, were identified using the clean_rawdata plugin (http://sccn.ucsd.edu/wiki/Plugin_list_process). Channels were rejected if they at any point during the recording flatlined for more than 5 seconds, if the channel correlated less than 0.85 with a reconstruction of this channel based on the surrounding electrodes, or if the channel had line noise relative to its signal that was four standard deviations or greater than the total channel population. Bad channels were interpolated from surrounding electrodes using EEGLab’s spherical interpolation, based on the method developed by Perrin, Pernier, Bertrand, and Echallier (1989). The data were segmented into stimulus-locked 1600-ms epochs, starting 200 ms before the target stimulus. Epochs were baseline-corrected relative to the time window 100-200 ms before the target stimulus. Eye blink activity was identified and removed using EEGLab’s default independent component analysis (ICA) method, which was followed by visual inspection and manual rejection of artifact components.

Group-level blind source separation was applied to estimate a single set of components that capture the representative activity from neural sources commonly expressed across the whole sample (Huster, Plis, & Calhoun, 2015). To do so, an equal number of trials were extracted from each participant. Only correct trials were used, with 81 congruent and 81 incongruent trials randomly selected for each subject. The group decomposition procedure requires the same number of trials per condition and subject, and thus the number of trials extracted per condition was set to keep the number of included trials high while at the same time excluding few subjects. In addition, the number of congruent trials was chosen to match the total number of incongruent trials to avoid biases in the estimation procedure. Then, group-SOBI (second-order blind identification) (Belouchrani, Abed-Meraim, Cardoso, & Moulines, 1997) was applied to the EEG, preceded by two consecutive principal component analysis steps (PCA; single subject and group level) as a means of data reduction (Eichele, Rachakonda, Brakedal, Eikeland, & Calhoun, 2011; Huster et al., 2015). First, for each participant, all trials were horizontally concatenated and z-standardized over channels, resulting in a two-dimensional matrix with electrodes as rows and time-points as columns spanning all 162 selected trials. This matrix was then entered into a subject specific PCA. Six PCs were selected from each participant, on average accounting for 70% of the variance in the single-subject data. 70% was chosen as a threshold because higher cutoffs resulted only in more noisy components without changing the main, stable components. These six components were extracted from each subject’s data, concatenated vertically (along the dimension originally representing channels), and entered into a second, group-level PCA. Again, six group-PCs were extracted and subjected to SOBI (with temporal lags up to 25 data points or 100 ms), resulting in six group-level component time-courses (Fig. 1). Finally, to examine between-subject differences in the group-level components, single-subject activations of those components were reconstructed by matrix multiplication of the original single-subject EEG time-courses with the demixing matrices resulting from the two preceding PCA’s and SOBI (see Eichele et al., 2011; Huster et al., 2015; Huster & Raud, 2018 for details). The matching of components to ERP peaks was based on component topographies, the degree of difference between incongruent and congruent trials, and the similarity in timing and appearance to the ERPs expected from the experimental paradigm. This led to the selection of two components (see below). Based on these criteria the peak activity in the difference wave of component 6 at 348ms was assumed to reflect a N2-like component. The peak of component 1 at 524 ms was assumed to reflect a P3-like component. In each of these components we subtracted the activity of congruent trials from that of incongruent trials and identified a peak of interest within this difference wave. At these peaks we extracted the area under the curve from a 40-ms time window surrounding each peak for use in further statistical analysis (hereafter referred to as time frames). From this point forward, the terms frontal negative component and parietal positive component will be used to describe the extracted component time frames. The summed area of the difference wave under the curve in each time frame will hereafter be referred to as that component’s strength. This measure of component strength is conceptually similar to regular difference scores in amplitude (delta amplitude). However, a component’s strength is in arbitrary units and the sign of each SOBI component peak is not indicative of the amplitude charge in the raw data before the decomposition. The actual amplitude of a component after its back projection to the EEG corresponds to the product of the component weight at a given electrode and its activity. For instance, if an electrode’s weight is negative, the corresponding EEG potential will be positive if the component activity is negative as well, whereas it will be negative if the component activity is positive.

**Figure 1.**
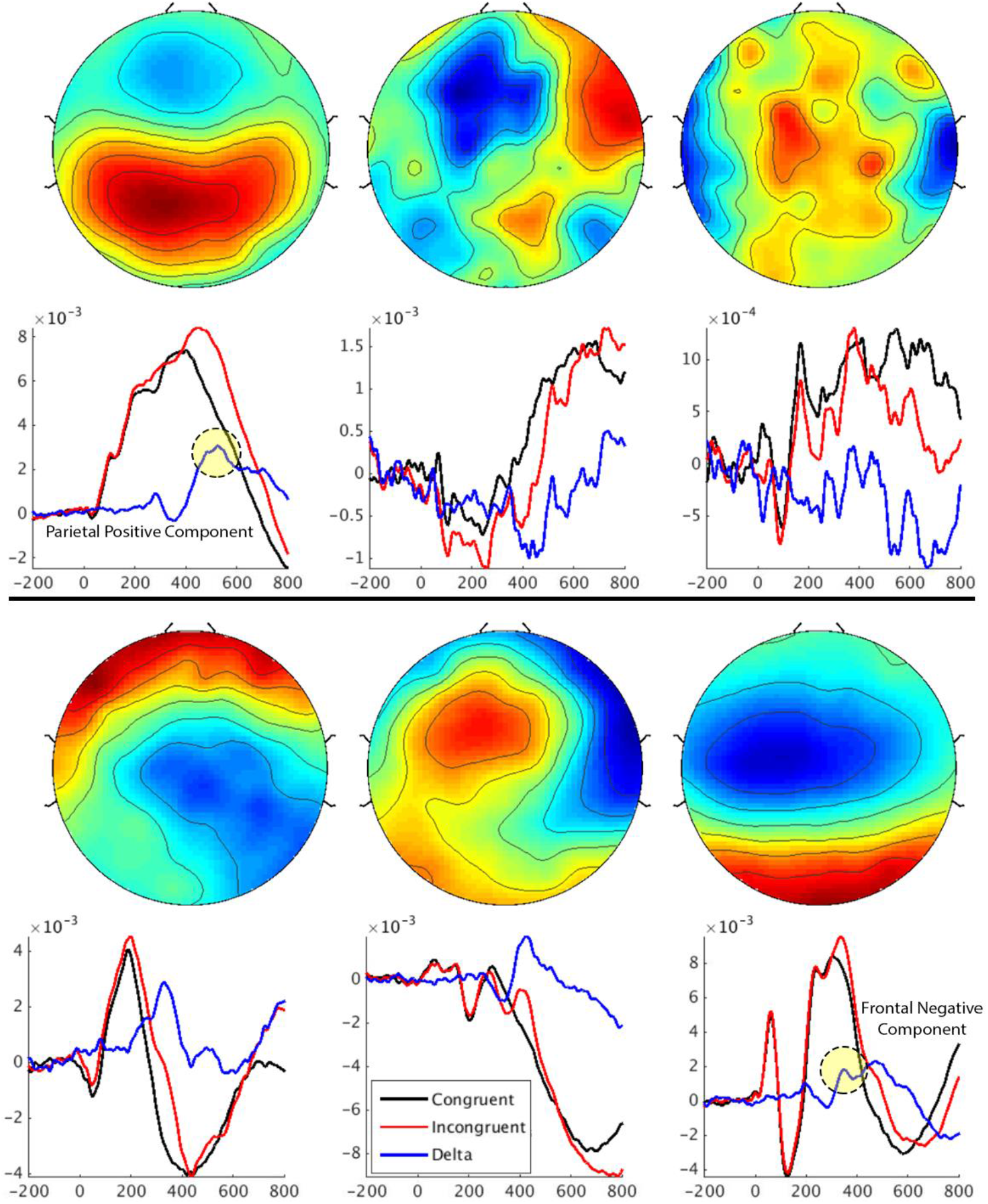
Component output of group-level SOBI decomposition of EEG epochs. Scalp topographies are shown above the associated component time courses. Time courses are sorted in congruent (black), incongruent (red) and delta (blue). Extracted peak deltas are marked by dotted circles for the parietal positive component (top left) and frontal negative component (bottom right).

### Statistical Analysis

Descriptive statistics and Pearson’s correlation analyses were used to characterize the sample and performance on the Flanker, Stroop and Anti-saccade tasks. Difference measures were calculated between incongruent and congruent events for the Flanker task for reaction time and accuracy for each subject, which were used as an index of congruency effect. Pearson’s correlation analyses were used to assess the relationship between these difference measures and age. Repeated measures ANOVAs with two factors with two levels each, congruency on the current trial and congruency on the preceding trial, were performed to test for post-conflict adjustments separately for reaction time and accuracy. For any ANOVA that revealed a significant interaction effect, follow-up T-tests were performed to determine the source of the interaction. Difference measures of reaction time and accuracy were calculated for event combinations of previous and current trial congruency where the T-tests revealed significant differences. Pearson’s correlations were calculated to assess the relationships between these difference measures and age. In all analyses that included EEG measures, only data from correct trials were used. Pearson’s correlations were used to assess the relationships between component strengths and age, while partial correlations, controlling for age, were used to assess the relationship between the strength of the EEG components and performance on the Flanker, Stroop and Anti-saccade tasks, as well as the congruency effect and Gratton effect difference measures.

## RESULTS

### Age-related Differences in Task Performance on the Flanker task

Reaction time on the Flanker task was negatively correlated with age for both congruent trials (r = −.61, p < .001) and incongruent trials (r = −.52, p < .001), indicating faster processing or responding with increasing age. Accuracy on the Flanker task was positively associated with age for congruent trials (r = .54, p < .001), but not for incongruent trials (r = .06, p = .517).

Pearson’s correlations between age and the difference score of incongruent and congruent reaction times (reaction time on incongruent trials minus reaction time on congruent trials) revealed a significant association (r = .24, p = .011), indicating a larger relative slowing, or congruency effect, for older participants. No significant relationship between age and the congruency effect was found for Flanker accuracy (r = −.17, p = .075).

Also on the Flanker task, repeated measures ANOVAs with two factors with two levels each, congruency on the current trial and congruency on the preceding trial, was performed to test for post-conflict adjustments (**Fig. 2**). The analysis of reaction times showed significant main effects of congruency on the current trial (F(1,107) = 1343.16, p < .001) and congruency on the preceding trial (F(1,107) = 19.75, p < .001), as well as an interaction between the two (F(1,107) = 20.43, p < .001). The same was found for accuracy, with significant main effects for congruency on the current trial (F(1,107) = 230.82, p < .001) and congruency on the preceding trial (F(1,107) = 119.81 , p < .001), and an interaction between the two (F(1,107) = 126.84, p < .001).

**Figure 2.**
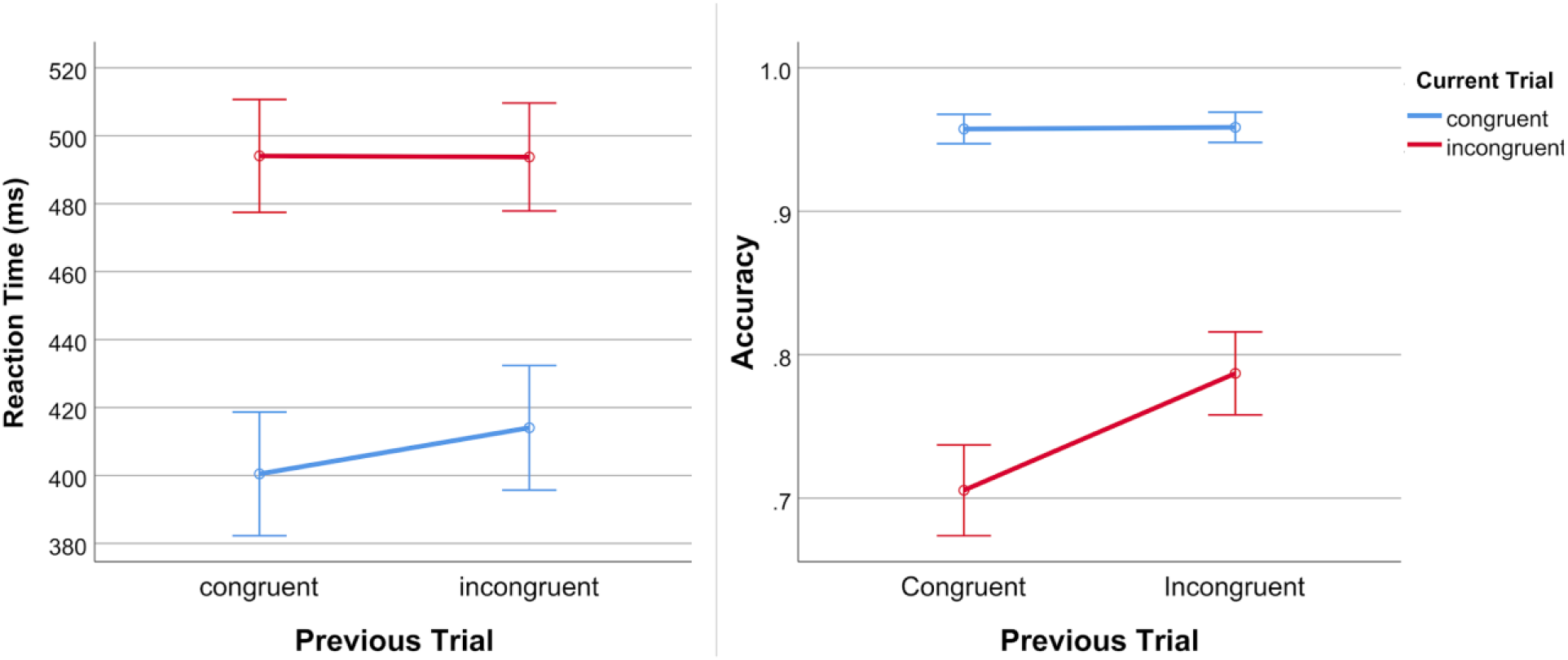
Line plots showing the interactions between current and previous trial type on reaction time (left) and accuracy (right).

Two paired samples t-tests were used to make post hoc comparisons between conditions on reaction time to clarify the source of the interaction effect. These revealed that reaction time was significantly faster for congruent events following congruent events (cC) (M=400.46, SD = 95.38) compared to congruent events following incongruent events (iC) (M=414.04, SD = 96.16) (t(108)= −6.96, p < .001), but no significant difference on reaction time between incongruent events following congruent events (cI) (M=494.07, SD = 87.09) and incongruent events following incongruent events (iI) (M=493.74, SD = 83.35) events (t(108)= .14, p = .89). Two paired-samples t-tests were used to make post-hoc comparisons between conditions on accuracy. These revealed that accuracy was significantly greater for iI (M=.79, SD = .15) events compared to cI (M=.71, SD = .17) events (t(108)= −11.87, p < .001), but that there was no significant difference on accuracy between cC (M=.96, SD = .05) and iC (M=.96, SD = .06) events (t(108)= −.45, p = .66). In sum, the results indicate a significant Gratton effect in our sample for both accuracy and reaction time, at least for some conditions, such that preceding incongruent trials increased reaction times on current congruent trials, and accuracy on current incongruent trials.

Difference measures were calculated for event combinations where the T-tests were significant (ΔC reaction time = cC reaction time – iC reaction time; ΔI accuracy = iI accuracy – cI accuracy). Of these, ΔI accuracy was significantly correlated with age (r = .25, p = .008), meaning that older participants had a greater relative improvement in accuracy on incongruent trials following incongruent trials compared to incongruent trials following congruent trials. We found no significant association between ΔC reaction time and age (r = .01, p = .923).

### Age-related Differences in Task Performance on the Stroop and Anti-saccade tasks

Completion time on the incongruent Stroop color-word interference task (with the congruent color naming condition regressed out) showed no significant correlation with age (r = −.12, p = .218). The Stroop ratio measure was however significantly correlated with age (r = −.44, p < .001), indicating that older participants had a smaller Stroop effect. Accuracy on the Anti-saccade task had a strong positive correlation with age (r = .68, p < .001), indicating greater cognitive control with increasing age.

### Associations Between Electrophysiological Components and Age

The strength of the parietal positive component showed a significant positive association with age (r = .24, p = .019), while the strength of the frontal negative component was not significantly correlated with age (r = .04, p = .720) (**Fig. 3**).

**Figure 3.**
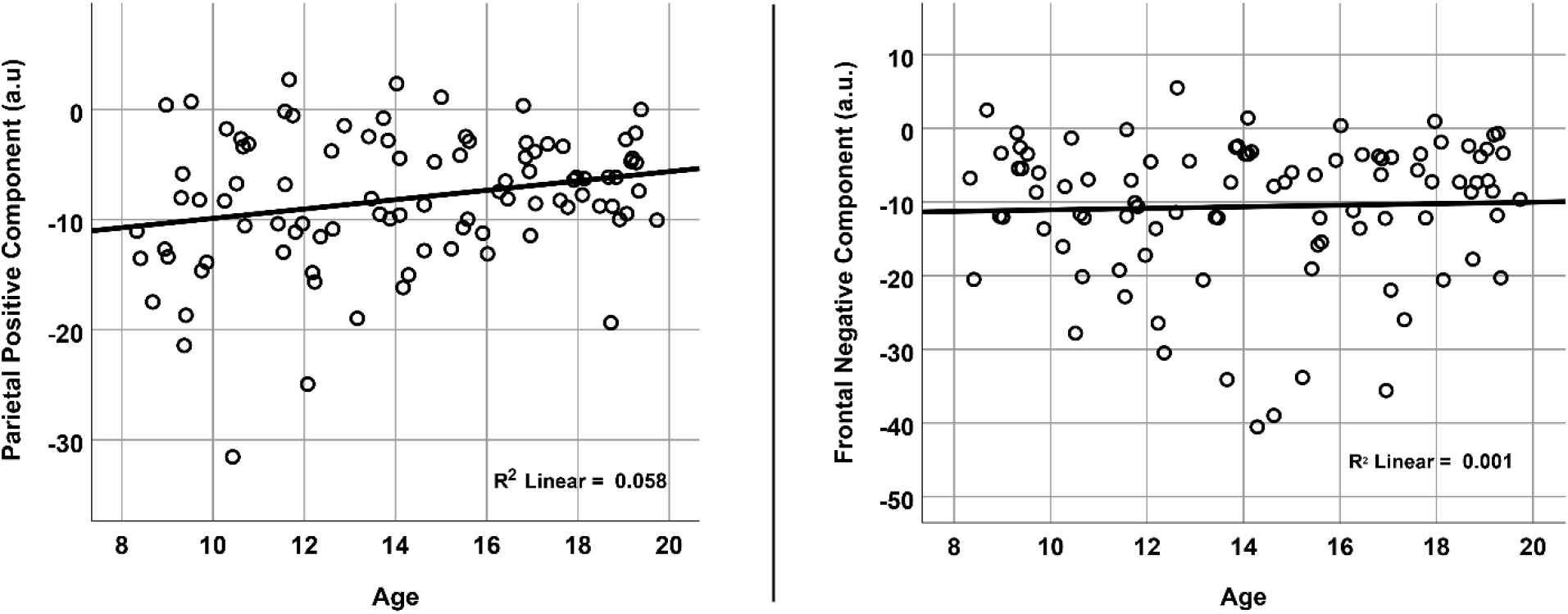
Scatter plots depicting relationships between age and component strength for the extracted time windows of each component. Signal strength is plotted in arbitrary units, and the signs of these values do not necessarily reflect the charge of the EEG amplitudes in the data before decomposition.

### Associations Between Electrophysiological Components and Task Performance

Partial correlations, controlling for age, revealed that neither the strength of the frontal negative component nor the parietal positive component were significantly associated with either accuracy or reaction time on the Flanker task. Strength of the frontal negative component was, however, independently of age, positively associated with performance on the Stroop task (r = .33, p = .001), meaning that a stronger frontal negative component was associated with faster task completion (with completion time on the Color naming condition regressed out). The same was found for the Stroop ratio measure (r = .38, p <.001). Strength of the frontal negative component also showed an age-independent negative association with Anti-saccade accuracy (r = −.31, p = .003), meaning that a stronger component was associated with higher accuracy. Strength of the parietal positive component was not significantly correlated with the Stroop task performance (Color naming regressed out: r = −.17, p = .127; ratio: r = − .14, p = .208) or Anti-saccade accuracy (r = .17, p = .109).

### Associations Between Electrophysiological Components and Congruency and Gratton Effects

Partial correlations, controlling for age, revealed a significant association between the strength of the positive parietal component and congruency effect on the Flanker task for reaction time (r = .217, p = .037), but not for accuracy (r = −.037, p = .723), indicating a greater relative reaction time on incongruent compared to congruent trials for participants with a stronger positive parietal component. No significant relationship was found between strength of the negative frontal component and the congruency effect for either reaction time (r = −.07, p = .504) or accuracy (r = .05, p = .619). Partial correlations, controlling for age, did not reveal any significant associations between the Gratton effect difference measures or any of the electrophysiological components.

## DISCUSSION

In this study, we investigated age-related differences in electrophysiological and behavioral indices of cognitive conflict processing across adolescence. Group-level blind source separation identified two electrophysiological components of interest from EEG data recorded during performance of a Flanker task: a frontal negative component corresponding to the N2, and a parietal positive component corresponding to the P3. A stronger negative frontal component was age-independently associated with better performance on Stroop and Anti-saccade tasks (overall reaction time, congruency RT, and accuracy). The N2-component also did not change in amplitude across age. The P3-component, on the other hand, was associated with performance monitoring processes such that, across subjects, stronger component activity coincided with more pronounced response slowing in incongruent relative to congruent trials. The strength of the positive parietal component also increased with age.

We further found evidence for congruency effects in trial sequences (aka Gratton effects) in the Flanker task for both reaction times and accuracies, but these effects were not associated with age.

### Relationships between age and task performance

For the Flanker task, older adolescents showed faster reaction times compared with younger participants. Task accuracy was positively associated with age for congruent, but not incongruent trials. On the Stroop Color-Word interference Task, we controlled for average reaction time on the color-naming task, creating a purer measure of interference control. Performance on this Stroop measure did not reveal any age-related differences. These results are the same as the ones reported by Lamm et al. (2006), but differ from what was found in several other Stroop studies (Ikeda, Okuzumi, & Kokubun, 2013; Ikeda et al., 2011; Prencipe et al., 2011), although these have used ratio measures instead of regression to control for the effect of color-naming performance. When we used a ratio measure similar to those used in these earlier studies, we did indeed find that older participants had a smaller Stroop effect. As ratio measures do not properly control for the denominator variable (Atchley et al., 1976; Lamm et al., 2006), it is not entirely clear if interference control as measured via the Stroop task improves substantially through adolescence or if a seeming improvement is explained by the overlapping development of other cognitive abilities. Performance on the Anti-saccade task was positively associated with age, in accordance with previous findings (Fukushima et al., 2000; Klein & Foerster, 2001; Luna et al., 2004). This indicates an improvement in interference control, but since we did not collect eye-tracking data as is commonly done for this task (Guitton, Buchtel, & Douglas, 1985), the difference in scores is difficult to disentangle from more general abilities of attention. Also, while all three tasks used involve cognitive control and resisting interference from irrelevant information, they likely load on a heterogeneous array of latent cognitive abilities. In the Miyake-Friedman model of executive control, the Stroop and Anti-Saccade task load on the same factor of response inhibition, while the Flanker task loads on a partially separate factor of Resistance to Distractor Interference (Friedman & Miyake, 2004; Miyake et al., 2000).

### Congruency and Gratton effects

In the Flanker task, the congruency effect on reaction time was greater for older participants, who showed a greater relative slowing on incongruent trials compared to congruent trials. This differs from the results of Erb and Marcovitch (2018), who found the congruency effect on accuracy and reaction time to decrease from childhood to adulthood. Based on the Gratton effect we should expect reaction time to be faster and accuracy to improve on trials in which the congruency is repeated compared to where congruency differs between trials (Gratton et al., 1992). Specifically, accuracy was greater on incongruent trials following incongruent trials compared to those following congruent trials. Also, reaction time was faster on congruent trials following congruent trials compared to those following incongruent trials. For accuracy, the size of the Gratton effect differed across age, with higher age being associated with a greater relative improvement in accuracy on incongruent trials following incongruent trials compared to those following congruent trials. This is again the opposite of what was reported by Erb and Marcovitch (2018). In contrast to their results, we found that both the congruency and Gratton effects seem play an increasing role in the task performance through development. The reason for these discrepant findings is unclear. Erb and Marcovitch (2018) argue that a reduced Gratton effect reflects a reduced influence from interference and might partly explain developmental improvements in interference control. However, as we also see a general improvement in task performance it could equally be argued that the increased Gratton effect in our sample reflects more efficient trial-wise adaptations.

### Relationships between age and electrophysiological components

The strength of the frontal negative component was not associated with age in our sample. This differs from what has been found for the N2 in children, where amplitude has generally been found to weaken with age (Lo, 2018), This might indicate that the N2 amplitude stabilizes in late childhood or early adolescence, although a longitudinal study by Isbell et al. (2018) indicates it might even reach stability in childhood. This contrasts to what has been found in the larger literature on the development of the ERN, an ERP with a neural origin and timing similar to the N2 (Gruendler et al., 2011; Van Noordt, Campopiano, & Segalowitz, 2016). The ERN seems to show continued development through adolescence (Davies, Segalowitz, & Gavin, 2004a, 2004b; Ladouceur, Dahl, & Carter, 2004; Santesso & Segalowitz, 2008; Santesso, Segalowitz, & Schmidt, 2006; Taylor, Visser, Fueggle, Bellgrove, & Fox, 2018). This is also what we found in the sample of the present study (Overbye et al., 2019). The dissociation in development between the N2 and ERN amplitude suggests that these components might have a greater functional difference than has been suggested in some earlier studies (Gruendler et al., 2011; Van Noordt et al., 2016).

The strength of the parietal positive component increased with age in our sample, becoming more positive. The positive association with age observed in our sample is the same relationship that has generally been described for the P3b in the literature (van Dinteren et al., 2014a, 2014b), including our own study on the same sample using an oddball paradigm (Overbye et al., 2018). This, along with its late timing and centro-parietal scalp topography suggests that the positive parietal component most resembles the P3b (Polich, 2007).

### Relationships between electrophysiological components and task performance

A stronger frontal negative component was negatively associated with improved task performance of both the Stroop task and the Anti-saccade task, independently of age. Such a relationship between N2 power and cognitive control function is the opposite of what was reported by Lamm et al. (2006), who found a stronger N2 to be associated with weaker performance on Stroop and the Iowa Gambling Task in adolescents. However, it is the same as has been reported before in adults (Huster et al., 2014). Surprisingly, while we found N2 strength to relate to these other measures of interference control, no such association was found for results on the Flanker task during which the EEG was recorded. These results may support the notion that the N2 reflects central aspects of cognitive control (Jodo & Kayama, 1992; Lezak, 1982), although the specifics of this relationship in terms of more narrowly defined functions is less clear. The strength of the parietal positive component was not associated with task performance on any of the cognitive tests or the Gratton effect, which could suggest that the processes indexed by the conflict induced P3b do not have a major influence on interference control. A stronger parietal positive component was however associated with a greater relative slowing on incongruent trials on the Flanker task. One interpretation of these results is that while the frontal negative component is related to global task performance, such as accuracy and overall reaction times, the parietal positivity is tied to more immediate trial-wise adaptations.

Seen together, our results on the development of interference control can be interpreted in different ways, as the improvements seen might relate to processes other than interference control. Differences between adolescents in interference control seem to be partially indexed by the N2, which seems to remain stable through adolescence. Differences between adolescents in N2 amplitude and interference control seem to be rooted in stable individual differences rather than developmental differences, although longitudinal studies are needed to confirm this.

## Conclusion

The present cross-sectional results suggest that the electrophysiological conflict-related frontal negative component does not change through adolescence, while the parietal positive component grows stronger. Greater strength of the frontal negative component was also found to be age-independently associated with better interference control. Higher age was associated with conflict having a larger impact, with a greater relative slowing of responses to conflict inducing stimuli. A Gratton effect was observed for our sample, with accuracy increasing and reaction time decreasing for Flanker trials of the same congruency relative to trial sequences that did not repeat. Age was also associated with a larger Gratton effect for accuracy. Together, the results portray a multifaceted and continued development of neurocognitive mechanisms for conflict processing across adolescence.

## Acknowledgments

This study was supported by the Department of Psychology, University of Oslo (to KO), and the Research Council of Norway (#230345, #288083, #223273).

## Conflict of Interest

The authors declare that they have no conflict of interest.

## Notes

### Competing Interest Statement

The authors have declared no competing interest.

### Summary of Updates

Changed distribution/reuse options to CC BY-NC

## REFERENCES

Atchley, W. R., Gaskins, C. T., & Anderson, D. (1976). Statistical properties of ratios. I. Empirical results. Systematic Zoology, 25(2), 137–148.

Belouchrani, A., Abed-Meraim, K., Cardoso, J.-F., & Moulines, E. (1997). A blind source separation technique using second-order statistics. IEEE Transactions on signal processing, 45(2), 434–444.

Bocquillon, P., Bourriez, J.-L., Palmero-Soler, E., Betrouni, N., Houdayer, E., Derambure, P., & Dujardin, K. (2011). Use of swLORETA to localize the cortical sources of target-and distracter-elicited P300 components. Clinical Neurophysiology, 722(10), 1991–2002.

Botvinick, M. M., Braver, T. S., Barch, D. M., Carter, C. S., & Cohen, J. D. (2001). Conflict monitoring and cognitive control. Psychological review, 108(3), 624–652.

Cavanagh, J. F., & Frank, M. J. (2014). Frontal theta as a mechanism for cognitive control. Trends in cognitive sciences, 18(8), 414–421.

Crone, E. A., & Steinbeis, N. (2017). Neural perspectives on cognitive control development during childhood and adolescence. Trends in cognitive sciences, 21(3), 205–215.

Davies, P. L., Segalowitz, S. J., & Gavin, W. J. (2004a). Development of error-monitoring event-related potentials in adolescents. Annals of the New York Academy of Sciences, 1021(1), 324–328.

Davies, P. L., Segalowitz, S. J., & Gavin, W. J. (2004b). Development of response-monitoring ERPs in 7-to 25-year-olds. Developmental neuropsychology, 25(3), 355–376.

Dehaene, S., Sergent, C., & Changeux, J.-P. (2003). A neuronal network model linking subjective reports and objective physiological data during conscious perception. Proceedings of the National Academy of Sciences, 100(14), 8520–8525.

Delis, D. C., Kaplan, E., & Kramer, J. H. (2001). Delis-Kaplan executive function system: Technical manual: Psychological Corporation.

Downes, M., Bathelt, J., & De Haan, M. (2017). Event-related potential measures of executive functioning from preschool to adolescence. Developmental Medicine & Child Neurology, 59(6), 581–590.

Egner, T. (2008). Multiple conflict-driven control mechanisms in the human brain. Trends in cognitive sciences, 72(10), 374–380.

Eichele, T., Rachakonda, S., Brakedal, B., Eikeland, R., & Calhoun, V. D. (2011). EEGIFT: group independent component analysis for event-related EEG data. Computational intelligence and neuroscience, 2011.

Erb, C. D., & Marcovitch, S. (2018). Deconstructing the Gratton effect: Targeting dissociable trial sequence effects in children, pre-adolescents, and adults. Cognition, 179, 150–162.

Erb, C. D., Moher, J., Song, J. H., & Sobel, D. M. (2018). Reach tracking reveals dissociable processes underlying inhibitory control in 5-to 10-year-olds and adults. Developmental science, 21(2), e12523.

Eriksen, B. A., & Eriksen, C. W. (1974). Effects of noise letters upon the identification of a target letter in a nonsearch task. Perception & psychophysics, 16(1), 143–149.

Folstein, J. R., & Van Petten, C. (2008). Influence of cognitive control and mismatch on the N2 component of the ERP: a review. Psychophysiology, 45(1), 152–170.

Friedman, D. (2003). Cognition and aging: a highly selective overview of event-related potential (ERP) data. Journal of Clinical and Experimental Neuropsychology, 25(5), 702–720.

Friedman, N. P., & Miyake, A. (2004). The relations among inhibition and interference control functions: a latent-variable analysis. Journal of Experimental Psychology: General, 133(1), 101.

Fukushima, J., Hatta, T., & Fukushima, K. (2000). Development of voluntary control of saccadic eye movements: I. Age-related changes in normal children. Brain and Development, 22(3), 173–180.

Gratton, G., Coles, M. G., & Donchin, E. (1992). Optimizing the use of information: strategic control of activation of responses. Journal of Experimental Psychology: General, 121(4), 480.

Gratton, G., Cooper, P., Fabiani, M., Carter, C. S., & Karayanidis, F. (2018). Dynamics of cognitive control: Theoretical bases, paradigms, and a view for the future. Psychophysiology, 55(3), e13016.

Gruendler, T. O., Ullsperger, M., & Huster, R. J. (2011). Event-related potential correlates of performance-monitoring in a lateralized time-estimation task. PloS one, 6(10), e25591.

Guitton, D., Buchtel, H. A., & Douglas, R. (1985). Frontal lobe lesions in man cause difficulties in suppressing reflexive glances and in generating goal-directed saccades. Experimental brain research, 58(3), 455–472.

Halgren, E., Baudena, P., Clarke, J. M., Heit, G., Liégeois, C., Chauvel, P., & Musolino, A. (1995). Intracerebral potentials to rare target and distractor auditory and visual stimuli. I. Superior temporal plane and parietal lobe. Clinical Neurophysiology, 94(3), 191–220.

Huster, R., Plis, S. M., & Calhoun, V. D. (2015). Group-level component analyses of EEG: validation and evaluation. Frontiers in neuroscience, 9, 254.

Huster, R., Westerhausen, R., Pantev, C., & Konrad, C. (2010). The role of the cingulate cortex as neural generator of the N200 and P300 in a tactile response inhibition task. Human brain mapping, 31(8), 1260–1271.

Huster, R. J., Enriquez-Geppert, S., Lavallee, C. F., Falkenstein, M., & Herrmann, C. S. (2013). Electroencephalography of response inhibition tasks: functional networks and cognitive contributions. International journal of psychophysiology, 87(3), 217–233.

Huster, R. J., Enriquez-Geppert, S., Pantev, C., & Bruchmann, M. (2014). Variations in midcingulate morphology are related to ERP indices of cognitive control. Brain Structure and Function, 219(1), 49–60.

Huster, R. J., & Raud, L. (2018). A tutorial review on multi-subject decomposition of EEG. Brain topography, 31(1), 3–16.

Ikeda, Y., Okuzumi, H., & Kokubun, M. (2013). Stroop/reverse-Stroop interference in typical development and its relation to symptoms of ADHD. Research in developmental disabilities, 34(8), 2391–2398.

Ikeda, Y., Okuzumi, H., Kokubun, M., & Haishi, K. (2011). Age-related trends of interference control in school-age children and young adults in the Stroop color-word test. Psychological reports, 108(2), 577–584.

Isbell, E., Calkins, S. D., Cole, V. T., Swingler, M. M., & Leerkes, E. M. (2018). Longitudinal associations between conflict monitoring and emergent academic skills: An event-related potentials study. Developmental Psychobiology.

Jodo, E., & Kayama, Y. (1992). Relation of a negative ERP component to response inhibition in a Go/No-go task. Clinical Neurophysiology, 82(6), 477–482.

Klein, C., & Foerster, F. (2001). Development of prosaccade and antisaccade task performance in participants aged 6 to 26 years. Psychophysiology, 38(2), 179–189.

Ladouceur, C. D., Dahl, R. E., & Carter, C. S. (2004). ERP correlates of action monitoring in adolescence. Annals of the New York Academy of Sciences, 1021(1), 329–336.

Lamm, C., Zelazo, P. D., & Lewis, M. D. (2006). Neural correlates of cognitive control in childhood and adolescence: Disentangling the contributions of age and executive function. Neuropsychologia, 44(11), 2139–2148.

Larson, M. J., Clayson, P. E., & Clawson, A. (2014). Making sense of all the conflict: a theoretical review and critique of conflict-related ERPs. International journal of psychophysiology, 93(3), 283–297.

Lezak, M. D. (1982). The problem of assessing executive functions. International journal of Psychology, 17(1-4), 281–297.

Lo, S. L. (2018). A meta-analytic review of the event-related potentials (ERN and N2) in childhood and adolescence: Providing a developmental perspective on the conflict monitoring theory. Developmental Review.

Luna, B., Garver, K. E., Urban, T. A., Lazar, N. A., & Sweeney, J. A. (2004). Maturation of cognitive processes from late childhood to adulthood. Child development, 75(5), 1357–1372.

Mahajan, Y., & McArthur, G. (2015). Maturation of mismatch negativity and P3a response across adolescence. Neuroscience letters, 587, 102–106.

McDermott, T. J., Wiesman, A. I., Proskovec, A. L., Heinrichs-Graham, E., & Wilson, T. W. (2017). Spatiotemporal oscillatory dynamics of visual selective attention during a flanker task. Neuroimage, 156, 277–285.

Miller, E. K., & Cohen, J. D. (2001). An integrative theory of prefrontal cortex function. Annual review of neuroscience, 24(1), 167–202.

Miyake, A., Friedman, N. P., Emerson, M. J., Witzki, A. H., Howerter, A., & Wager, T. D. (2000). The unity and diversity of executive functions and their contributions to complex “frontal lobe” tasks: A latent variable analysis. Cognitive psychology, 41(1), 49–100.

Østby, Y., Tamnes, C. K., Fjell, A. M., Westlye, L. T., Due-Tønnessen, P., & Walhovd, K. B. (2009). Heterogeneity in subcortical brain development: a structural magnetic resonance imaging study of brain maturation from 8 to 30 years. Journal of Neuroscience, 29(38), 11772–11782.

Overbye, K., Huster, R. J., Walhovd, K. B., Fjell, A. M., & Tamnes, C. K. (2018). Development of the P300 from childhood to adulthood: a multimodal EEG and MRI study. Brain Structure and Function, 223(9), 4337–4349.

Overbye, K., Walhovd, K. B., Paus, T., Fjell, A. M., Huster, R. J., & Tamnes, C. K. (2019). Error processing in the adolescent brain: Age-related differences in electrophysiology, behavioral adaptation, and brain morphology. Developmental cognitive neuroscience, 38, 100665.

Perrin, F., Pernier, J., Bertrand, O., & Echallier, J. F. (1989). Spherical splines for scalp potential and current density mapping. Electroencephalogr Clin Neurophysiol, 72(2), 184–187.

Polich, J. (2007). Updating P300: an integrative theory of P3a and P3b. Clinical Neurophysiology, 118(10), 2128–2148.

Prencipe, A., Kesek, A., Cohen, J., Lamm, C., Lewis, M. D., & Zelazo, P. D. (2011). Development of hot and cool executive function during the transition to adolescence. Journal of experimental child psychology, 108(3), 621–637.

Roberts, R. J., Hager, L. D., & Heron, C. (1994). Prefrontal cognitive processes: Working memory and inhibition in the antisaccade task. Journal of Experimental Psychology: General, 123(4), 374.

Santesso, D. L., & Segalowitz, S. J. (2008). Developmental differences in error-related ERPs in middle-to late-adolescent males. Developmental Psychology, 44(1), 205–217.

Santesso, D. L., Segalowitz, S. J., & Schmidt, L. A. (2006). Error-related electrocortical responses in 10-year-old children and young adults. Developmental science, 9(5), 473–481.

Segalowitz, S., & Davies, P. L. (2004). Charting the maturation of the frontal lobe: an electrophysiological strategy. Brain and cognition, 55(1), 116–133.

Smith, M. E., Halgren, E., Sokolik, M., Baudena, P., Musolino, A., Liegeois-Chauvel, C., & Chauvel, P. (1990). The intracranial topography of the P3 event-related potential elicited during auditory oddball. Electroencephalography and clinical neurophysiology, 76(3), 235–248.

Stroop, J. R. (1935). Studies of interference in serial verbal reactions. Journal of experimental psychology, 18(6), 643.

Tamnes, C. K., Fjell, A. M., Westlye, L. T., Ostby, Y., & Walhovd, K. B. (2012). Becoming consistent: developmental reductions in intraindividual variability in reaction time are related to white matter integrity. The Journal of Neuroscience, 32(3), 972–982.

Tamnes, C. K., Ostby, Y., Fjell, A. M., Westlye, L. T., Due-Tonnessen, P., & Walhovd, K. B. (2010a). Brain maturation in adolescence and young adulthood: regional age-related changes in cortical thickness and white matter volume and microstructure. Cerebral cortex, 20(3), 534–548.

Tamnes, C. K., Østby, Y., Walhovd, K. B., Westlye, L. T., Due-Tønnessen, P., & Fjell, A. M. (2010b). Neuroanatomical correlates of executive functions in children and adolescents: a magnetic resonance imaging (MRI) study of cortical thickness. Neuropsychologia, 48(9), 2496–2508.

Taylor, J. B., Visser, T. A., Fueggle, S. N., Bellgrove, M. A., & Fox, A. M. (2018). The error-related negativity (ERN) is an electrophysiological marker of motor impulsiveness on the Barratt Impulsiveness Scale (BIS-11) during adolescence. Developmental cognitive neuroscience, 30, 77–86.

van Dinteren, R., Arns, M., Jongsma, M. L., & Kessels, R. P. (2014a). Combined frontal and parietal P300 amplitudes indicate compensated cognitive processing across the lifespan. Frontiers in aging neuroscience, 6, 294.

van Dinteren, R., Arns, M., Jongsma, M. L., & Kessels, R. P. (2014b). P300 development across the lifespan: a systematic review and meta-analysis. PloS one, 9(2), e87347.

Van Noordt, S. J., Campopiano, A., & Segalowitz, S. J. (2016). A functional classification of medial frontal negativity ERPs: Theta oscillations and single subject effects. Psychophysiology, 53(9), 1317–1334.

Wechsler, D. (1999). Manual for the Wechsler Abbreviated Intelligence Scale (WASI) The Psychological Corporation. San Antonio, Tx.

Wronka, E., Kaiser, J., & Coenen, A. M. (2012). Neural generators of the auditory evoked potential components P3a and P3b. Acta Neurobiologiae Experimentalis, 72(1), 51–64.

Zelazo, P. D., & Müller, U. (2002). Executive function in typical and atypical development. Blackwell handbook of childhood cognitive development, 445–469.

